# A science friction story – Molecular interactions in semiflexible polymer networks

**DOI:** 10.1101/2021.09.22.461249

**Authors:** Paul Mollenkopf, Dusan Prascevic, Martin Glaser, David M. Smith, Jörg Schnauß

## Abstract

Established model theories, developed to capture the mechanical behavior of soft complex materials composed of semiflexible polymers assume entropic interactions between filaments to determine the mechanical response. In recent studies, the general accepted tube model has been challenged in terms of its basic assumption about filament-filament interactions, but also because of its predictions regarding the frequency dependence of the elastic modulus in the intermediate frequency regime. A central question is how molecular interactions and friction between network constituents influence the rheological response of isotropic entangled networks of semiflexible polymers. It was shown that friction forces between aligned pairs of actin filaments are not negligible. Here, we systematically investigate the influence of friction forces and attractive interactions on network rheology by means of a targeted surface modification. We show that these forces have a qualitative and quantitative influence on the viscoelastic properties of semiflexible polymer networks and contribute to the response to nonlinear deformations. By comparing two polymer model systems with respect to their surface compositions we give a possible explanation about the origin of acting forces on a molecular level.

## 1. Introduction

Semiflexible polymers are intensively studied materials due to their special mechanical properties and their significant role in complex soft matter. Biological systems rely on semiflexible polymers as building blocks of intracellular scaffolds and extracellular matrices. Protruding as an essential component for cell shape regulation and force generation to promote cell migration and division, they take on a special role within the intracellular skeleton^1^. Semiflexibility implies a persistence length in the range of the polymers contour length (*L*_*p*_ ≈ *L*)^2^.

Actin as the most prominent representative^2,4–6^ has been subject to numerous in vitro studies^7–12^. Accompanied by the experimental characterization, theoretical models have been developed to describe the emergence of viscoelastic properties in semiflexible polymer networks^13–15^. The characterization of individual filaments is theoretically embedded within the generally accepted wormlike chain (WLC) model^16–18^. Based on the WLC, the tube concept is constructed to capture network properties of entangled polymer solutions. It exclusively accounts for entropic interactions between filaments neglecting any attractive or repulsive effects caused by the molecular structure. Based on these assumptions, fluctuation modes are suppressed due to surrounding polymers establishing a confinement in a tube-like region^15,19–21^. Some important signatures in the frequency dependent rheology are qualitatively reflected by the model. The drop of the elastic modulus G’ towards low frequencies is associated with disengagement of filaments through reptation. The crossover point where the loss modulus transcends the elastic modulus denotes the transition from low frequency network properties dominated by filament interactions to single filament dominated high frequency behavior^15^. Both of these experimentally measurable features are explained by the model^22^.

However, experimental studies on filamentous actin (F-actin) and DNA based semiflexible polymers questioned the model’s central scaling prediction with respect to the persistence length^23,24^. Additionally, recent work emphasized the importance of attractive interfilamentous interactions in networks caused by hydrophobicity or electrostatic interactions, which are not considered by the tube model^25,26^. Despite representing some crucial effects in the viscoelastic network behavior, the tube model predicts a flat plateau of G’ in the intermediate frequency range. Experimental data, however, have consistently revealed a weak power law behavior, which is related to filament-filament interactions^25–29^.

Employing optical tweezers, Ward *et al* showed surprisingly high friction forces between a pair of aligned actin filaments^30^. The friction between two filaments was found to depend on polarity as well as on depletion force induced cohesiveness and scaled logarithmically with the sliding velocity^30^. In multifilament bundles, contractile forces that can be attributed to pairwise filament interactions determine the relaxation behavior of bundles^31^. The mechanical response of actively excited actin bundles formed by depletion forces suggests that interactions and crosslinking are difficult to distinguish^31,32^. On a network level recent studies found that attractive filament-filament interactions have a qualitative and quantitative impact on the rheological behavior of isotropic networks of semiflexible polymers ^25,26,29,33^. A theory that accounts for attractive sticky interactions in networks and explains the experimentally observed power law behavior was proposed by Kroy and Glaser^34^. The so-called glassy wormlike chain (GWLC) extends the wormlike chain by incorporating interactions between individual filaments in a network by presuming a glassy surrounding^34^. This is mathematically described by the stretching of the WLCs long wavelengths Eigenmodes with an Arrhenius-like exponential factor. The model has been successfully employed in experimental research providing a quantitative characterization of non-specific interactions between networks components^25,26,29^. However, the interactions that are investigated and quantified with the model are not specified and their origins remain speculative. Ward *et al* established an approach to reduce the friction between actin filaments by a surface modification with polyethylene glycol (PEG) molecules that act as an entropic spring to prevent friction interactions in an anisotropic arranged two-filament system ^30^. We adapted this procedure to the prerequisites for networks of two different polymer model systems. By comparison of their rheological properties, we isolated the impact of friction and/or attractive forces on network mechanics. We compared F-actin to a DNA-based polymer models system (DX5 nanotubes) built according to construction plans of DNA nanofabrication^35^. The DX5 nanotubes, introduced by Ekani-Nkodo *et al*^36^, exhibit similar mechanical properties as F-actin^37^. With comparable contour lengths and bending rigidities of respective individual filaments, the DX5 nanotubes differ to actin in their surface composition with respect to topography, hydrophobicity and electric charges, each of which is plausible to contribute to friction and/or attractive filament-filament interactions. Previous studies compared different polymer model systems with regard to their “stickiness” to understand their network architectures and their role within the cytoskeleton^25,26,29^. We systematically investigated the influence of surface-surface interactions between filaments in isotropic networks, pioneering the exploration of molecular interactions in multi-filament systems and their intrinsic impact on higher ordered structures.

## 2. Results

### 2.1. Linear network rheology

To isolate the impact of friction forces and attractive inter-filamentous interactions on the mechanics of isotropic network structures, we systematically investigated the viscoelastic properties of F-actin as well as DNA nanotubes. We compared each system to its modified versions. We synthesized three different modifications of actin as well as DX 5 tubes by decorating the filaments’ surfaces with PEG molecules of three different lengths. The success of the modifications was checked via gel electrophoresis (Fig 2, ESI†). We found bands indicating a successful PEG binding to the respective surfaces. PEG modification is an established method to reduce nonspecific binding of quantum dots in live cell assays^38^. Here, the surface coating prevents attractive filament-filament interactions on length scales depending on the end-to-end distance of the PEG chain, which is ∼6.35 *nm* per kDa in outstretched configuration^39^. With this approach, we were able to systematically alter mutual inter-filament interactions^40^.

**Figure 1.**
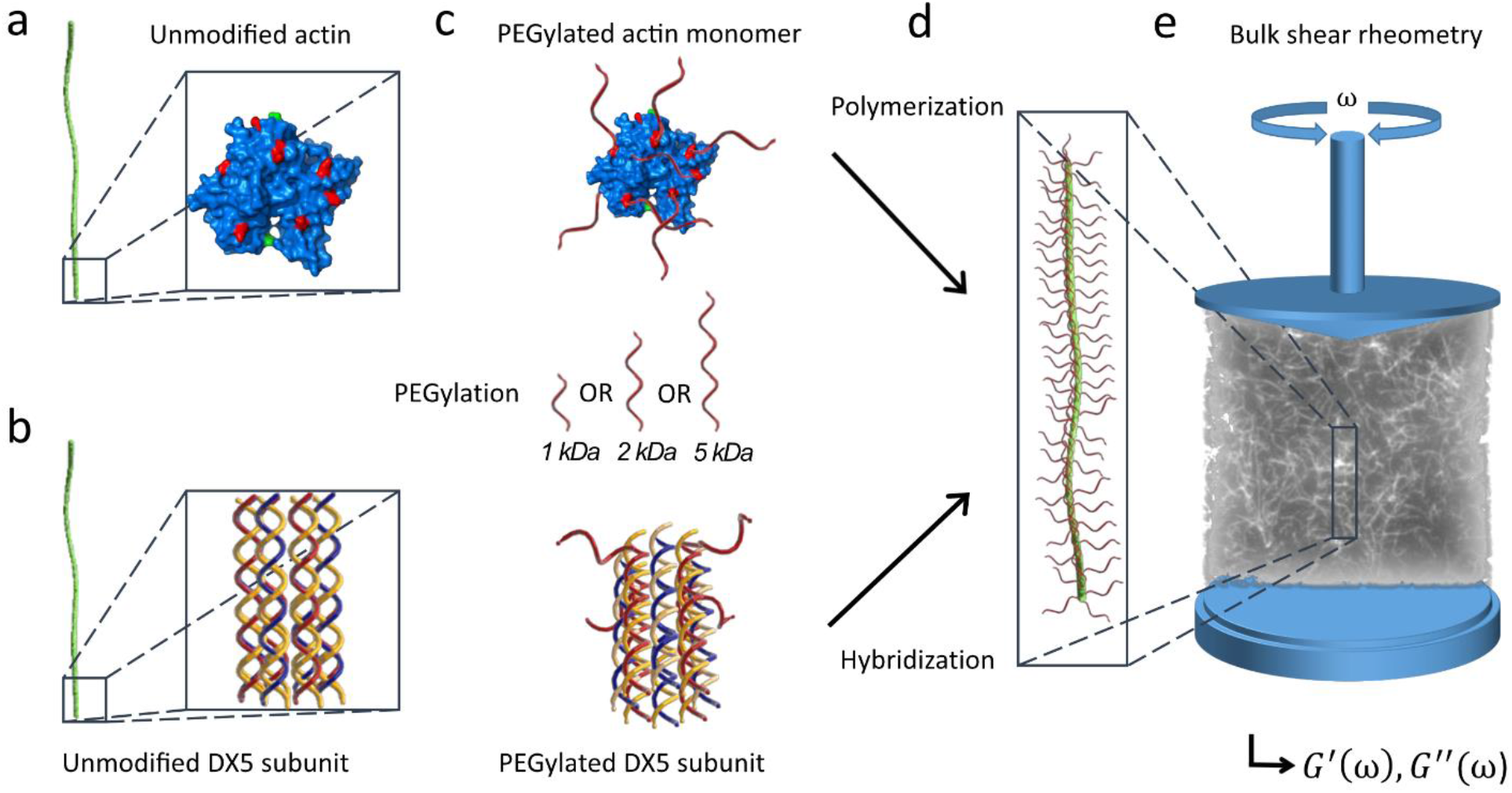
Modification of semiflexible filaments with PEG molecules. Actin monomers (a) have primary amines like lysine exposed to the molecules surface which serve as target for a modification via NHS Ester reaction chemistry (Polymerization relevant domains are labeled green^3^). Semiflexible polymers can be artificially recreated via DNA nanotechnology. DX5 nanotubes are a DNA based reference system, composed of 5 different single DNA strands which hybridize in unit elements (b) that build up to elongated tubes and can be modified via copper-free click chemistry. PEG modified unit elements / monomers (c) hybridize / polymerize into filaments with PEGylated surfaces. PEG molecules act as spacers to screen filament-filament interactions over three controlled distances determined by the length of the respective attached PEG molecules (d). The viscoelastic properties of reconstituted networks are experimentally accessible via bulk shear rheometry (e).

**Figure 2.**
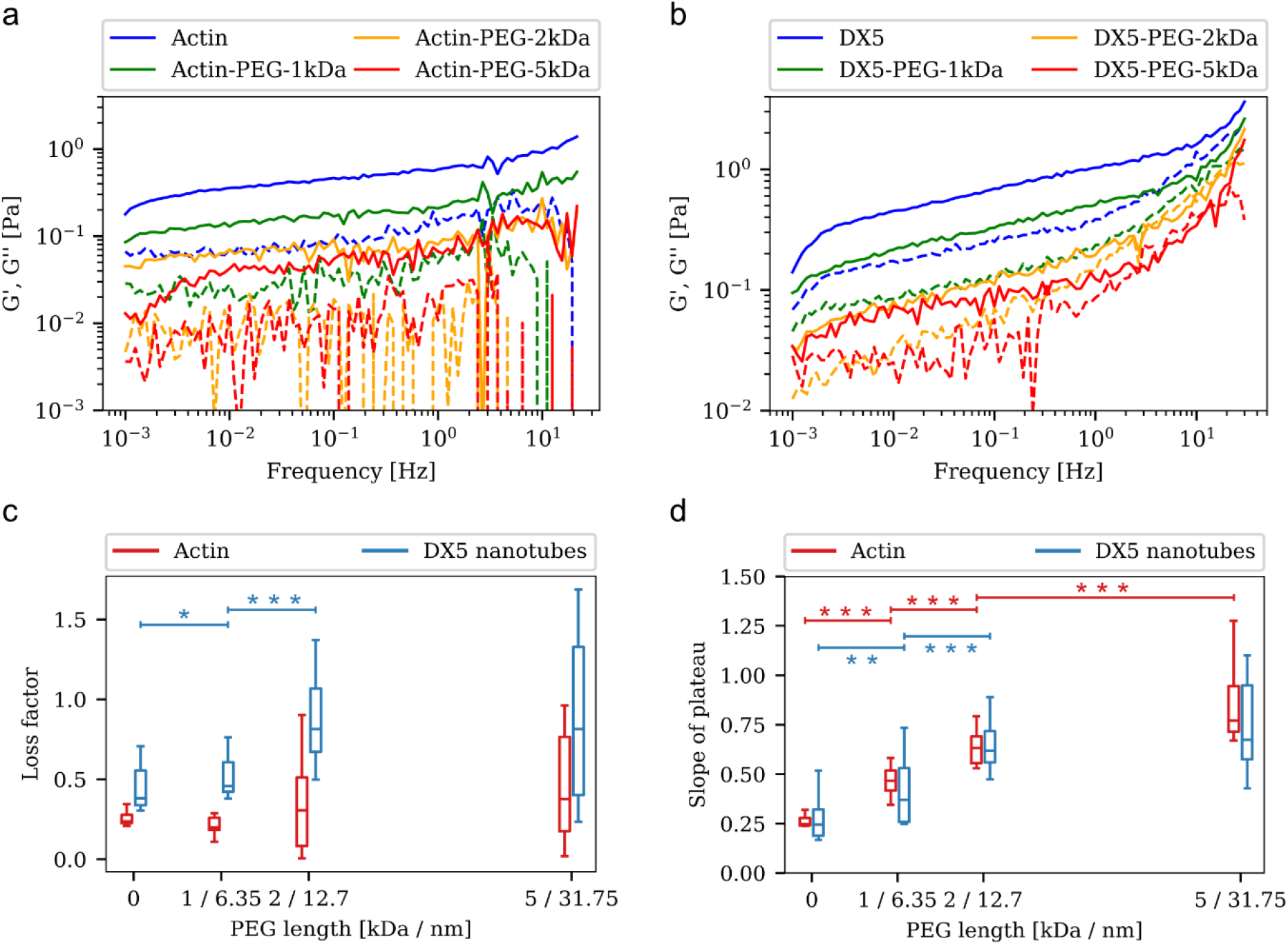
The complex shear modulus *G*^∗^ decreases for increasing lengths of PEG molecules attached to the respective filament surfaces (a, b). The loss factor 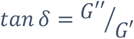 exhibits a general increase upon PEG modification, indicating a tendency towards viscous behavior. In contrast to the DNA-based reference system (DX5 nanotubes), we found a non-monotonic increase in *tan δ* for actin networks. This comes along with a general increase in contour length for modified actin filaments (Fig. 3 ESI†). A decreasing slope of the elastic modulus is associated with a transition to networks of highly inter-attractive networks and cross-linked systems, where the frequency dependence is less pronounced. We found increasing slopes for increasing spacers between interacting filaments, reflecting the successive elimination of the contribution of attractive interactions (d). Significant different distributions, according to a Mann*-*Whitney*-*U*-*Test, *are* indicated by * symbols with significance levels p < 0.05 (∗), p < 0.01 (∗∗) and p < 0.001 (∗∗∗).

Viscoelastic properties of suspended networks of semiflexible polymers are experimentally accessible via bulk shear rheology yielding the complex shear modulus G*(f) = G’(f) + i G’’(f). The results of linear rheology are summarized in Fig 2. Elastic moduli as well as loss moduli decreased with increasing lengths of PEG chains for both F-actin and DX5 nanotubes (Fig. 2 a,b). The loss factor tan *δ*, quantifying the relation between viscosity and elasticity as 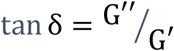, shows a general increase upon PEG modification. This reflects a gradual transition from elasticity to more viscosity dominated networks. In contrast to the DX5 system, F-actin networks showed a non-monotonic behavior in tan *δ*. Measurements suggested that actin filaments, in contrast to DX5 nanotubes can exhibit increased contour lengths when modified with PEG molecules (Fig. 3, ESI†). We suspect that the non-monotonic increase of loss factors is due to the fact that the increase in elasticity by longer actin filaments compensates for the fluidization of the networks. By fitting the power-law behavior of the elastic modulus over frequencies ranging from 0.1 to 10 Hz, we determined the slopes of the elastic plateau for both systems (Fig. 2 d). With 0.26, the slope value for unmodified F-actin networks is in agreement with previous findings^29^. The value for unmodified DX5 tube networks was calculated to be 0.28 and is thus comparable to F-actin networks. Both systems displayed slopes that incrementally increased up to values of 0.86 for actin-PEG-5kDa and 0.73 for DX5-PEG-5kDa. These findings confirm the results of previous experimental studies pointing to higher values for loss factor and slope of plateau for presumably less interactive polymer systems^25,26,29,41^.

**Figure 3.**
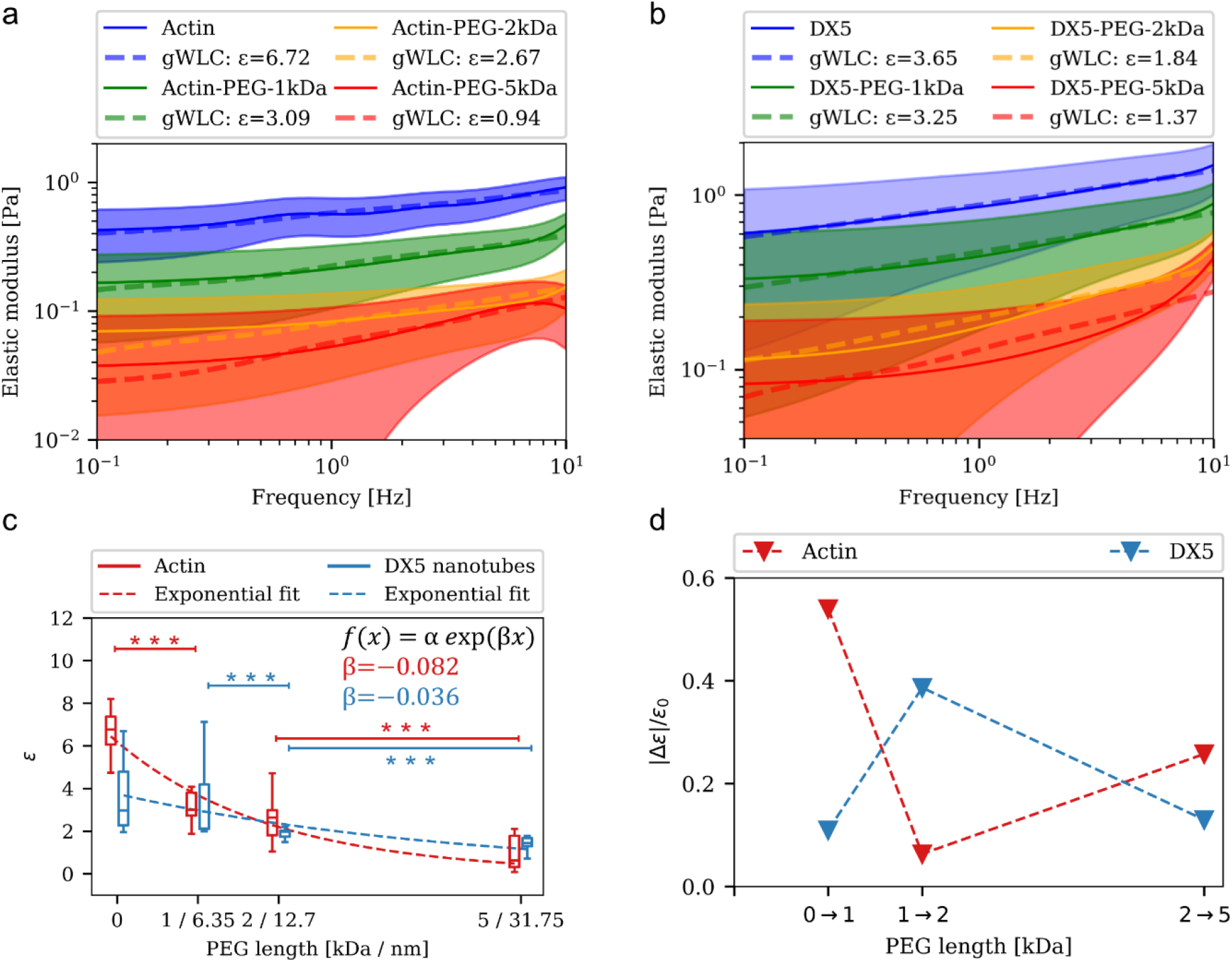
Attractive interactions between individual filaments in networks of entangled semiflexible polymers decrease in strength for increasing length of PEG chains covering the filaments’ surface. This is deduced from the frequency dependent elastic modulus for actin (a) and DX5 nanotubes (b). Solid lines show the mean curves of measured G’(f), smoothed with a univariate spline fit. Dashed lines represent the fits according to the GWLC model. Each curve represents the mean of 12 measurements. The ε values resulting from fitting to the GWLC model are listed in the legend and plotted with error bars representing respective standard deviations (c). Dashed lines represent exponential fits according to *f*(𝓍) = α exp(*β*𝓍). The resulting β values indicate a more drastic decrease for the F-actin networks. The relative gradual change upon stepwise extension of PEG molecules is illustrated in (d). Significant different distributions, according to a Mann*-*Whitney*-*U*-*Test, are indicated by * symbols with significance levels *p* < 0.05 (∗), *p* < 0.01 (∗∗) and *p* < 0.001 (∗∗∗).

### 2.2 Glassy wormlike chain

The impact of the surface modification on the frequency dependent viscoelastic properties of the investigated networks can be explained in the frame of the GWLC model^34^. The filaments’ mode relaxation times τ_λ_ > τ_*Λ*_ of all Eigenmodes of (half) wavelength λ longer than a characteristic interaction length Λ are stretched by the factor exp(εN). Here, 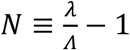 gives the interactions per wavelength λ and corresponds to the entanglement length^34^. Unspecific interactions between filaments in a network of entangled semiflexible polymers are governed in the stretching parameter ε, which determines the slowdown of the polymer dynamics and gives a quantification of the interaction strength^25,26,29,34^. A more detailed description of this model is presented in the ESI†.

The linear rheology data was analyzed by fitting the storage modulus G’(f) to the mean curves of the measured data with

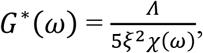

where ω = 2π*f* and χ(ω) is the micro-rheological response function of the GWLC to a point force at its ends in the high frequency limit^42^. The fitting routine was implemented in a self-written Python script. Contour lengths and persistence lengths for actin as well as DX5 nanotubes were measured via fluorescence microscopy and used accordingly for the evaluation (Fig.3 and Fig.4 ESI†). As we compare networks of same molarities, we emanate from similar mesh sizes, leaving us with the characteristic interaction length Λ and the stretching parameter ε as the only non-fixed fitting parameters. We used a mesh size of ∼1µ*m* for DX5 networks and ∼1.4µ*m* for F-actin networks. The drag coefficient was set to ζ _┴_ = 2 *mPa s*, a representative value for water.

We found ε = 6.72 for untreated actin and ε = 3.65 for DX5 nanotubes, respectively. We interpret the stretching parameter ε as a strength of attractive interactions between individual filaments in a network. The magnitude of this parameter is sensitive to small deviations in the preset of fixed parameters such as mesh size or contour lengths of filaments. Hence, we did not only consider absolute values but also accounted for the gradual decline of the strength of interactions upon surface modification.

We examined the stretching parameter ε for both polymer systems and their respective modifications. Distributions differed significantly for all combinations within the F-actin derivatives, except for actin-PEG-1kDa compared to actin-PEG-2kDa. For the DX5 nanotube system, the only significant reduction between directly neighbored distributions is recorded for the stepwise screening length increase from DX5-PEG-1kDa to DX5-PEG-2kDa. Interestingly, the gradually reduction of the interaction strength is not the same for actin and DX5 tube networks. The decrease of the interaction strength ε can be described with an exponential function for both systems (Figure 3 c). With an exponent of *β* = −0.082 for actin, the overall reduction is clearer than for the DX5 system, where we found an exponent of *β* = −0.036. The step-wise reduction, illustrated in Figure 3d, reveals a more pronounced interaction decrement for F-actin networks of filaments covered with 1 kDa PEG. Ward *et al* showed that friction forces between aligned actin filaments depend on their orientation of polarity with respect to each other, concluding that the surface topography itself has a strong impact on the inter-filament friction^30^. Actin filaments, in contrast to DX5 nanotubes, have a surface structure that promotes friction. In this context, the drastic reduction can be attributed to the compensation of steric forces, that are originating from surface-surface friction. With a diameter of approximately 7 nm^43^, the roughness of the actin filaments’ surface is compensated by 1kDa PEG molecules with a contour length of roughly 6.35 *nm*. With a further increase of PEG molecule lengths from 1 kDa to 2 kDa, ε reduces more for the DNA based system. Hydrophobic interactions between molecules reportedly decay exponentially like van der Waals dispersion forces over a range of ∼10 *nm* and are an order of magnitude stronger^44^. Monomeric actin consists of 375 amino acids, some of which are hydrophobic^45^. However, during protein folding, most of them are inherently turned to the monomers interior to prevent contact to its aqueous surrounding^46,47^. Consequently, the reduction due to the elimination of hydrophobic effects is presumably stronger for the DX5 nanotubes, wherein adenine and thymine in particular promote hydrophobic driven interactions^48^. Electrostatic interactions can be orders of magnitudes stronger than the van der Waals interactions^49^. Being relevant for DNA based materials as well as amino acids^50,51^, they presumably contribute to the filament-filament interactions of both systems in a comparable manner. The estimation of the influence of electrostatic forces is not trivial as they strongly depend on the ions present in the buffer solution that can compensate repulsive interactions between polymers^52^.

### 2.3 Nonlinear rheology

We tested the mechanical behavior of F-actin and DX5 nanotube networks under high deformations by applying a constant strain rate and measuring the stress responses. We exposed the samples to strains in the nonlinear regime prior to calculating the differential modulus K, defined as the local derivative of stress σ over strain γ as described by Semmrich *et al*^41^. Consistent with previous studies we found a weak stiffening behavior for F-actin networks^25,26,29,53^. This is reflected in a slight increase of K as illustrated in Fig. 4a. For modified F-actin networks as well as for DX5 nanotube networks and its modified versions (Fig. 4b) we found no stiffening behavior under nonlinear deformations. This is in line with the ε values calculated from the linear data. The yield stresses, which denote the transition from elastic to plastic behavior, shifted to lower values for decreasing associated ε values obtained from the model for the linear data (Fig.3). Network stiffening as a response to large deformations has been shown for various intermediate filaments and in particular for keratins, which are known to provide structure and protection against large deformations to epithelial cells^29,54,55^. F-actin networks exhibit weak strain stiffening which is increased by the presence of crosslinking complexes^41,56,57^. Previous studies showed that systems consisting of filaments which are assigned with high ε values express drastic stiffening upon shear deformationsa^25,26,29^. Here we show, that the stiffening behavior for F-actin networks can be eliminated by screening interactions on distances of few nanometers. From this we conclude that actin filaments in entangled networks interact strongly enough to influence the nonlinear network behavior.

**Figure 4.**
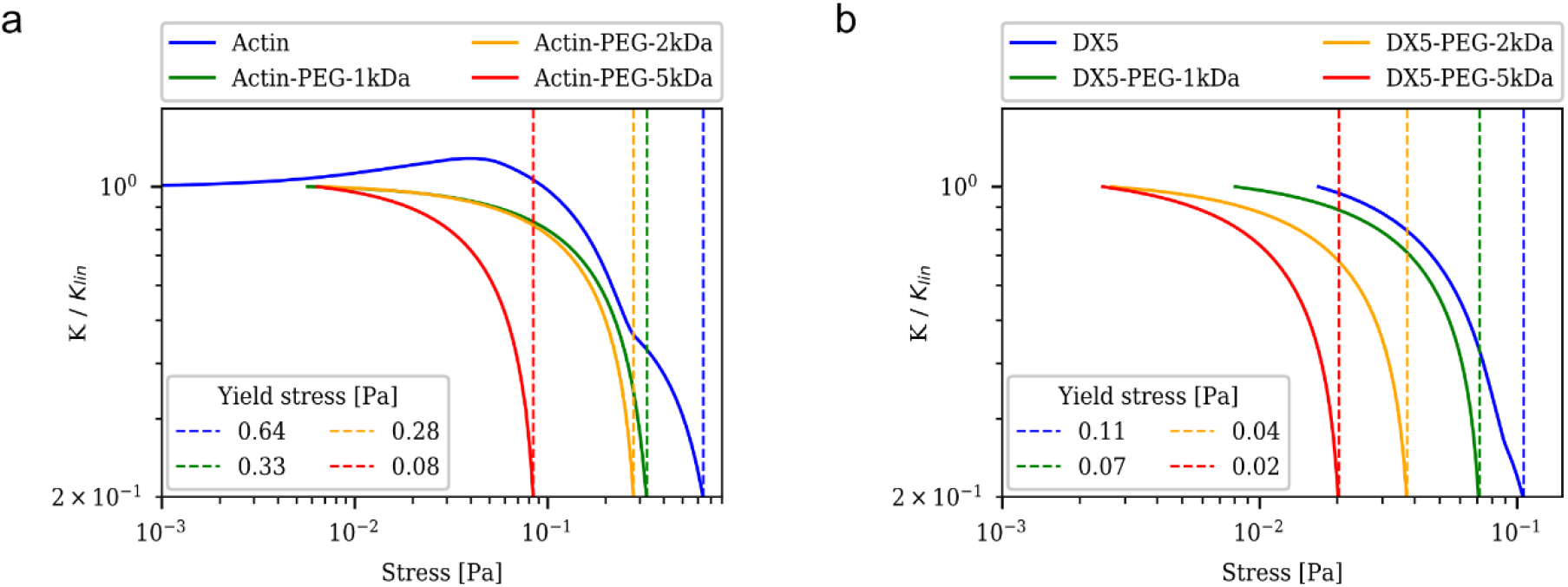
Attractive interactions affect the mechanic behavior under nonlinear network deformations. Stiffening is weak for actin networks and vanishes for networks of PEG modified actin filaments, manifesting in a flattening of the differential shear modulus. The DX5 nanotube networks exhibit no stiffening behavior. The yield stress is shifted to lower stress values for increasing lengths of PEG molecules for both investigated systems.

## 3. Discussion

Previous *in vitro* studies investigated the influence of cross-linkers of various kinds on the mechanics of isotropic networks of semiflexible polymers^2,12,56–59^. Here we show, that eliminating attractive interactions and friction forces has a comparable quantitative and qualitative effect on network rheology. By a systematic screening of surface-surface interactions we determined rheological signatures associated with friction or attractive interactions.

We found a progressive network softening for increasing spacer molecule lengths between filaments. This is accompanied by a significant general increase in the slopes of the elastic modulus over frequency and a transition to less elastic networks.

The evaluation with the GWLC model yielded gradually rising relaxation dynamics for long wavelength Eigenmodes indicating a decrease of the impact of attractive interactions to the overall network rheology. From this we conclude that there is a causal relationship between the apparent power-law behavior as well as the network softening with the dominance of mutual interactions of semiflexible polymers in an entangled network, confirming the assumptions of earlier studies^25,26,29^. These results are not explainable in the frame of the tube model which does not consider attractive interactions on a molecular level. The increase in power law behavior upon reduction of interactions contradicts the theory which predicts a flat plateau for the elastic modulus while not considering attractions between network constituents.

Mathematically explaining the slowdown of filament dynamics with unspecified interactions, the GWLC model does not provide an explanation about their fundamental nature. The decomposition of acting forces is impeded by the fact that the range alone is not a criterion to be able to differentiate explicitly between different types of interactions. However, we can provide a rough estimate of the various forces. Ward *et al* measured surprisingly high friction forces for a pair of actin filaments moved against each other^30^. In compliance with this, we assume that steric hindrance due to filament surface texture plays a similar role in entangled network rheology. On intermediate distances, our results show a stronger reduction of attraction for the DNA based system suggesting that hydrophobicity causes attractive forces. This is in line with recent findings that fibrils of α-synuclein stiffen significantly due to temperature dependent interactions between hydrophobic patches on the fibril surface^33^.

The described interactions and friction forces influence the rheological behavior of networks under large deformations in the nonlinear regime. Strain stiffening and strain softening are critical properties that determine a systems mechanical integrity. Our results suggest, that friction forces can induce a weak strain stiffening behavior in F-actin networks which play a crucial role for cellular mechanics^13,42,56,57,60^.

Mutual interactions between filaments have an impact on the mechanic of higher ordered structures. On a cellular level, hydrophobic forces between cytoskeletal filaments due to the presence of deuterium oxide are suspected to slow down cellular dynamics like proliferation and migration on a remarkable degree^11,61^. This highlights the relevance of interactions for cellular systems that mechanically rely on networks of semiflexible polymers.

## 4. Conclusions

The mechanical behavior of materials composed of semiflexible polymers strongly depends on interactions between filaments. The response of cross-linked system is tension-dominated and comparably easy to describe theoretically^22^. For an all-encompassing understanding of entangled network rheology, forces between molecules have to be comprised in theoretical models. Besides hydrophobicity and electrostatic interactions, the filaments’ surface structure in particular must be accounted for.

## Materials and Methods

### 1. Protein preparation, polymerization and PEGylation

G-actin was prepared from rabbit muscle and stored at -80 °C in G-Buffer (2 mM sodium phosphate buffer pH 7.3, 0.2 mM ATP, 0.1 mM CaCl2, 1 mM DTT, 0.01% NaN3) as described previously^62^. For experimental use, small volumes of monomeric actin were aliquot and thawed until needed. Fluorescently labeled actin was prepared by polymerizing G-actin at 5 mM in a 1:1 ratio with phalloidin–tetramethylrhodamine B isothiocyanate (phalloidin–TRITC – Sigma-Aldrich Co.). Polymerization of actin samples was induced by adding 1/10 volume fraction of 10 times concentrated F-buffer (20 mM sodium phosphate buffer pH 7.3, 1 M KCl, 10 mM MgCl2, 2 mM ATP, 10 mM DTT).

Actin filaments were covered with PEG (Polyethylenglycol) according to a self-elaborate conjugation protocol. After polymerizing in phosphate F-Buffer at a pH of 7.2 to 7.3 for 90 minutes, actin was incubated overnight with a 10-fold excess of NHS-Ester modified PEG (1kDa, 2kDa, 5kDa) that reacts with primary amines on the surface of actin monomers to yield stable amide bonds. The modification is applied to filamentous actin to prevent the occupation of polymerization relevant binding sites. To depolymerize actin and simultaneously remove unbound PEG from the solutions, samples were dialyzed in dialysis bags with 14kDa pore sizes against 1.5l phosphate G-Buffer.

### 2. DNA Crossover tube hybridization and PEGylation

The design as well as the necessary oligomers for the hybridization of the DNA nanotubes were adapted from Ekani-Nkodo *et al*.^36^ [Table SII]. Stock solutions were prepared by resuspending lyophilized DNA oligomers purchased from Biomers.net. The concentration of each stock solution was confirmed spectrophotometrically by a NanoDrop 1000 (Thermo Fisher Scientifc Inc., USA) at a wavelength of 260nm. DNA Nanotube networks were assembled at a concentration of 20 µM by mixing the required single stranded DNA sequences SE1 to SE5 in equimolar concentration in an assembly buffer containing 40mM Tris-acetate, 1mM EDTA and 12.5mM Mg 2+ (pH 8.3). These strands were hybridized in a Professional Standard PCR Thermocycler (Core Life Sciences Inc., USA) by denaturation for 10min at 90°C and complementary base pairing for 20h between 80° C and 20° C, lowering the temperature by 0.5K every 10min. The hybridization can be divided into two sub-processes. Unit elements which are hybridized at around 50°C stack up to tubular structures at distinctly lower temperatures of around 30°C. After hybridization the constituted nanotubes were stored at room temperature. For visualization the oligomer SE3 was modified with the fluorescent Cyanine dye 3 with two additional spacer thymine bases in between. DNA nanotubes were labeled by partially or fully replacing the unlabeled oligo SE3 by SE3-Cy3. DNA tubes were covered with PEG molecules according to a self-elaborate conjugation protocol. The oligomer SE1 was modified with a DBCO group and consequentially incubated with a 5-fold excess of PEG-azide (1kDa, 2kDa, 5kDa) overnight. Unbound PEG was removed either by an ethanol precipitation or by an Amicon filtration.

### 3. Rheology

Shear rheology measurements were performed with a strain controlled ARES rheometer (TA Instruments, USA) and a cone–plate geometry with a diameter of 25 mm and a gap width of 50 µm.

Actin/actin-PEG was polymerized between plate and cone for 2 hours at 20 °C after initiating polymerization by adding 10-fold F-buffer and water to a final volume of 175µl. To prevent interfacial elasticity artifacts, the cone was surrounded with F-buffer, eliminating air exposure for polymers. The sample chamber was sealed with a cap equipped with wet sponges to prevent evaporation. Polymerization was monitored with a dynamic time sweep by a measurement every 120 s at a frequency of 1 Hz and a strain of 5%. Only samples that were in equilibrium at the end of the time sweep were considered for further analysis. The linear regime was measured with a dynamic frequency sweep with a strain of 5% and 20 points per decade.

DNA nanotubes were put on the rheometer pre-hybridized as described above in the same sample volumes as for actin measurements. After a 120-minute equilibration phase, monitored by a dynamic time sweep with a measurement every 120 s at a frequency of 1 Hz and a strain of 5%, the same measurement protocol as for actin was applied.

A transient step rate test at a strain rate of 0.1 *s*^−1^ was used to measure the non-linear strain regime. The differential shear modulus K was determined from the resulting stress–strain curves with a self-written Python script. K was calculated as the gradient of the spline fit smoothed stress data divided by the strain step width. The linear value *K*_*lin*_ was defined at the first non-negative stress value. Negative stress values, particularly for small strains, appear due to measurement limitations as well as a result of the fitting routine.

