## Supplementary material for "A science friction story – Molecular interactions in semiflexible polymer networks": Supporting information.docx

–

| 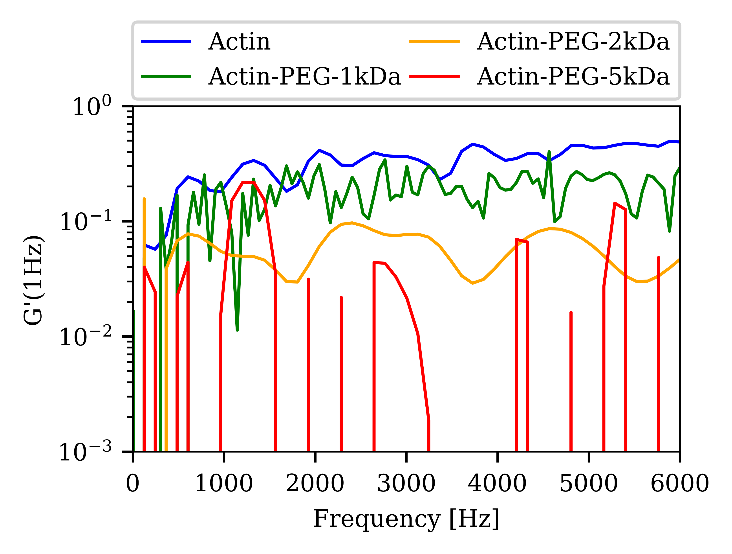**a** | 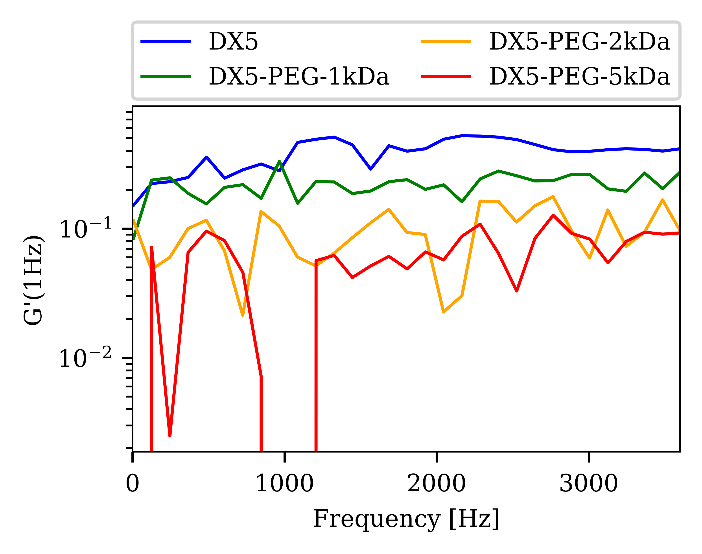**b** |
| --- | --- |
| 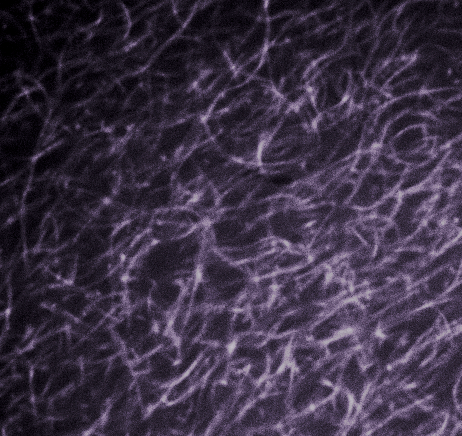**c** | 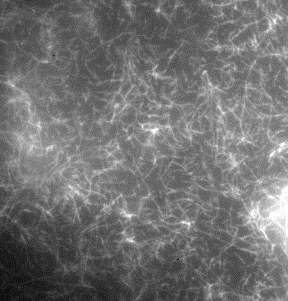**d** |

Figure 1 Modified actin monomers polymerize into networks of filamentous actin (d), equipped with an interaction hindering cover in the form of polyethylene glycol molecules attached to the surface via a NHS-Ester reaction. For network visualization we used a fluorescent PEG compound. Actin polymerization was monitored via rheometry where the elastic modulus at 1 Hz was measured every 2 minutes (a). DX5 nanotube networks (d) were hybridized prior to loading onto the rheometer. The equilibration was monitored by measuring the elastic modulus at 1 Hz.

### **Table A1**: DNA strands used for the DX5 nanotube system

| Strand | Sequence |
| --- | --- |
| SE1 | ctcagtggacagccgttctggagcgttggacgaaacT |
| SE2 | gtctggtagagcaccactgagaggta |
| SE3 | ccagaacggctgtggctaaacagtaaccgaagcaccaacgct |
| SE4 | cagacagtttcgtggtcatcgtacct |
| SE5 | cgatgacctgcttcggttactgtttagcctgctctac |
| SE1-PEG | ctcagtggacagccgttctggagcgttggacgaaacT-${PEG}_{n kDa}$ , $n$ =1,2,5 |

**Gel electrophoresis:**

| **a**  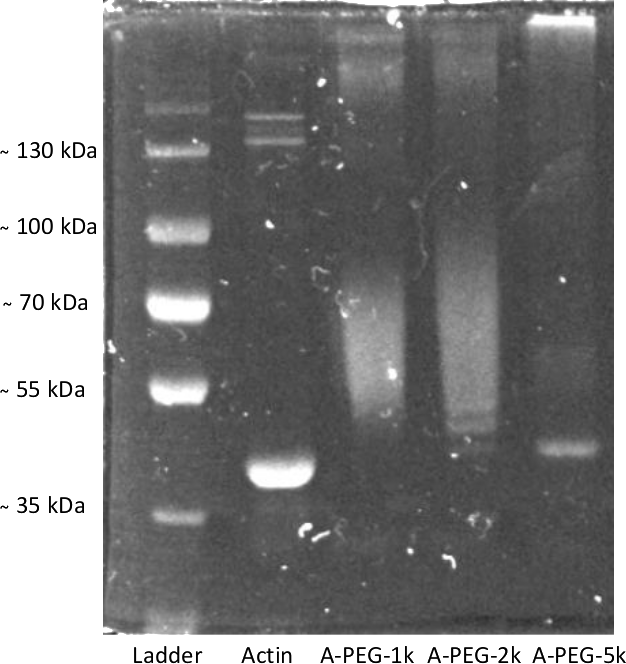 | **b**  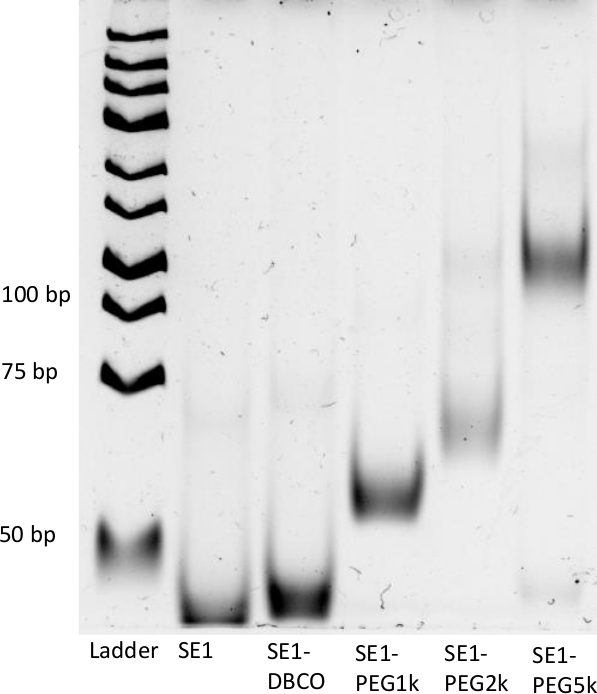 |
| --- | --- |

*Figure 2*: Gel electrophoresis was used to verify the success of applied PEG surface modifications. (a) A ThermoScientific PageRuler Plus was used as size standard to compare the size differences between monomeric actin and actin monomers decorated with polyethylene glycol chains of weights 1 kDa, 2 kDa and 5 kDa (from left to right). The actin band is at approximately 42 kDa. For the modification with PEG molecules we observe broad bands, indicating a distribution of multiple bond PEG molecules per actin molecule. For 5 kDa a thin band at 42 kDa indicates a proportion unmodified monomeric actin. A faint broad band between ̴50 kDa and 60 kDa indicates monomers with several 5 kDa PEG molecules. The bright band on top of the column shows, that many actin monomers are decorated with multiple PEG molecules. The modified actin monomers are too large to move through the gel. (b) A Thermo Sientific GeneRuler DNA ladder was used to compare the single stranded DNA sequence SE1 (see above) to its modified versions. To enable a copper free click chemistry reaction to bind azide-conjugated PEG to the sequence, SE1 was conjugated with DBCO prior to the PEG modification step. For the modification with 5 kDa we observe a faint band at the height of unmodified SE1, indicating unmodified strands like for actin.

**Contour lengths and persistence lengths actin & actin-PEG:**

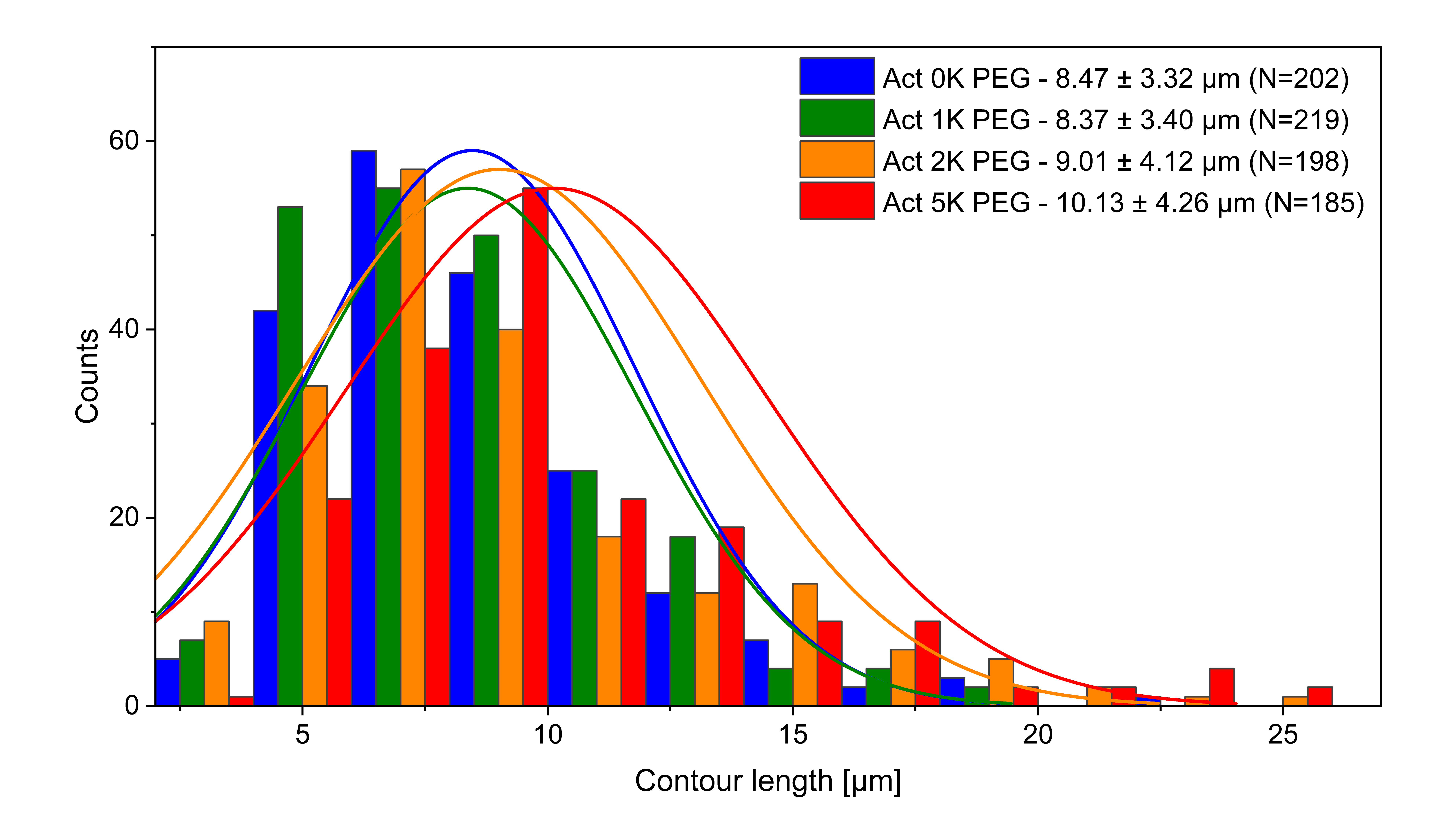

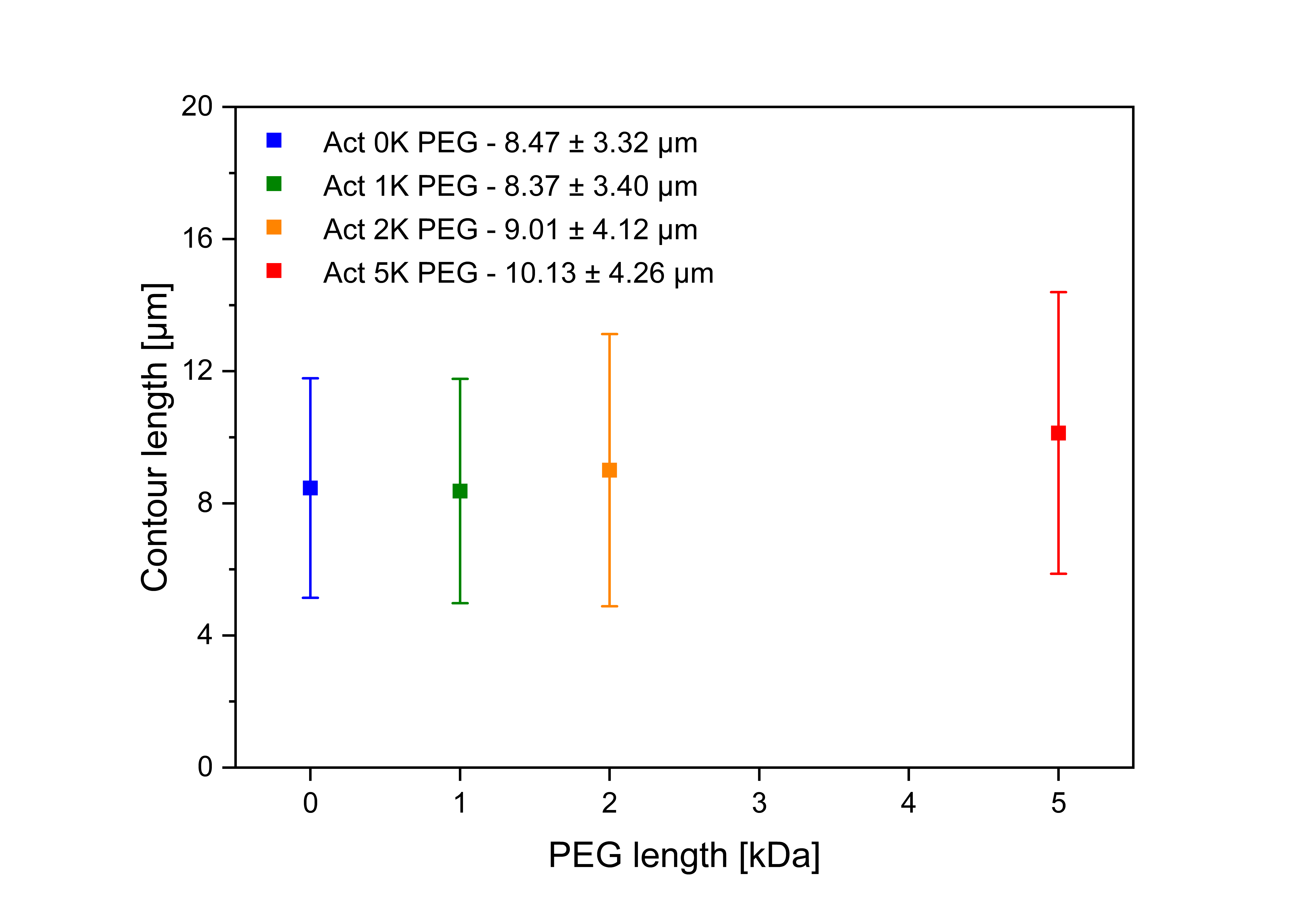

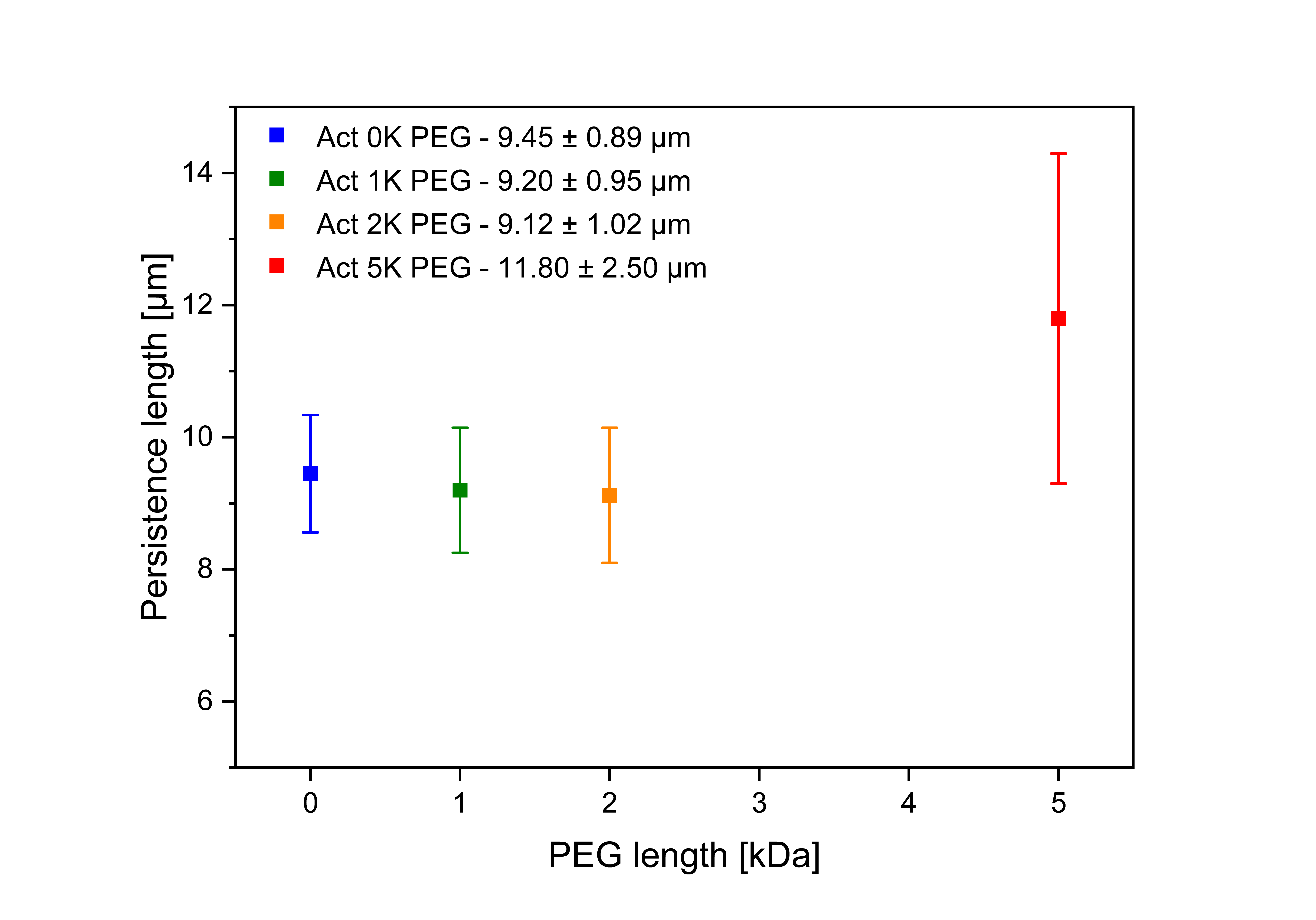

*Figure 3*: Contour lengths and persistence lengths for F-actin and the PEG modified versions of F-actin.

**Contour lengths and persistence lengths DX5 & DX5-PEG:**

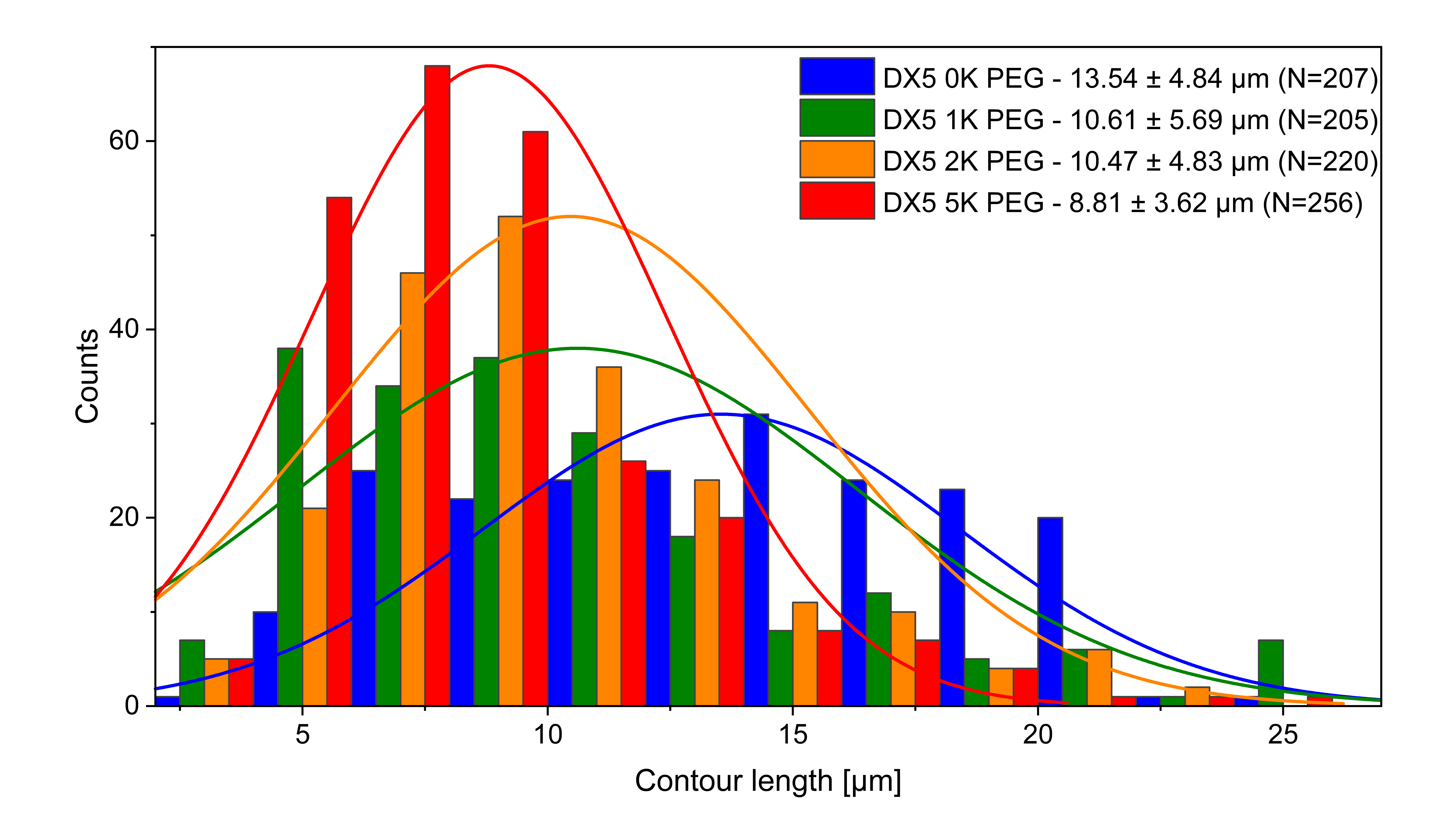

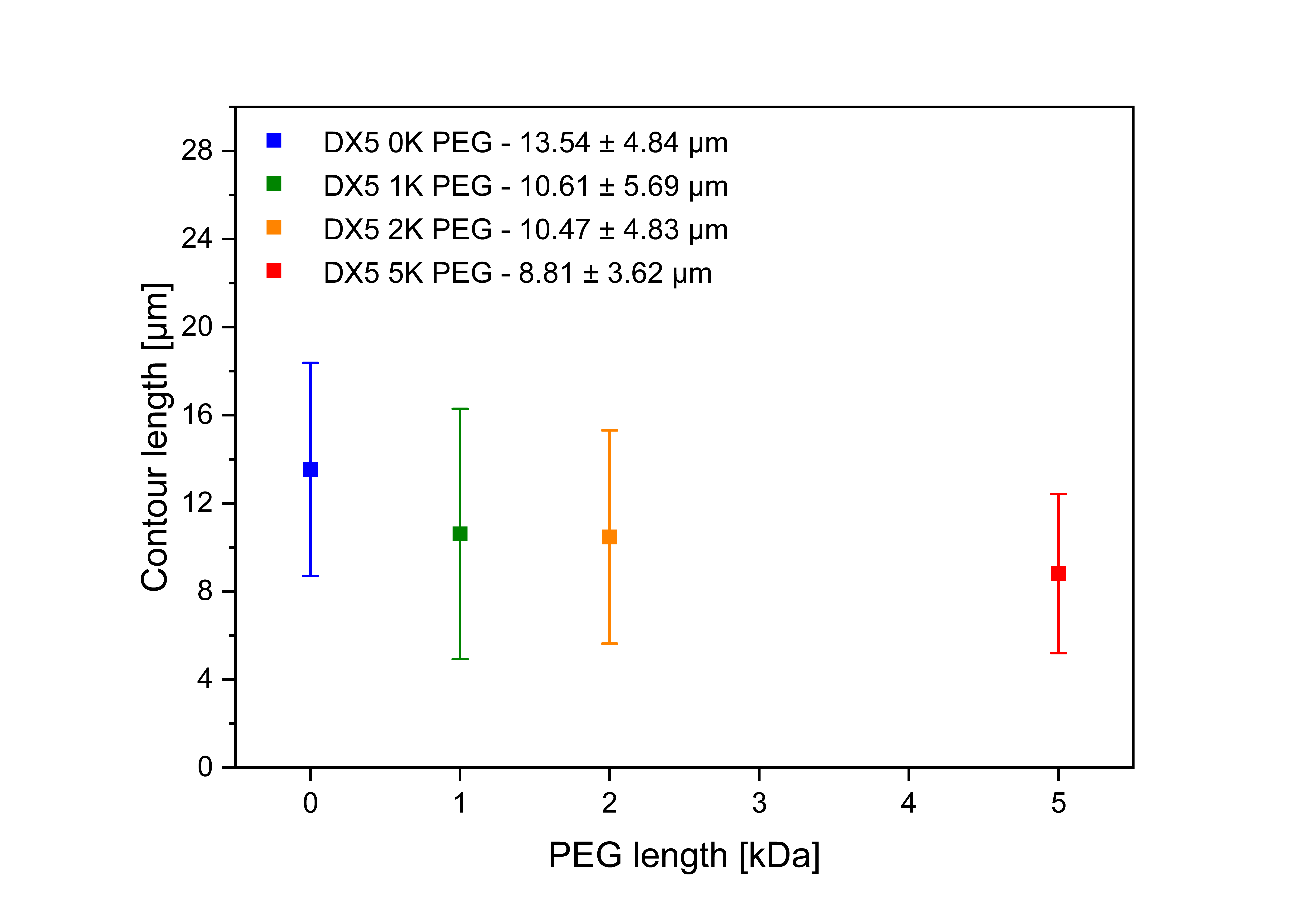

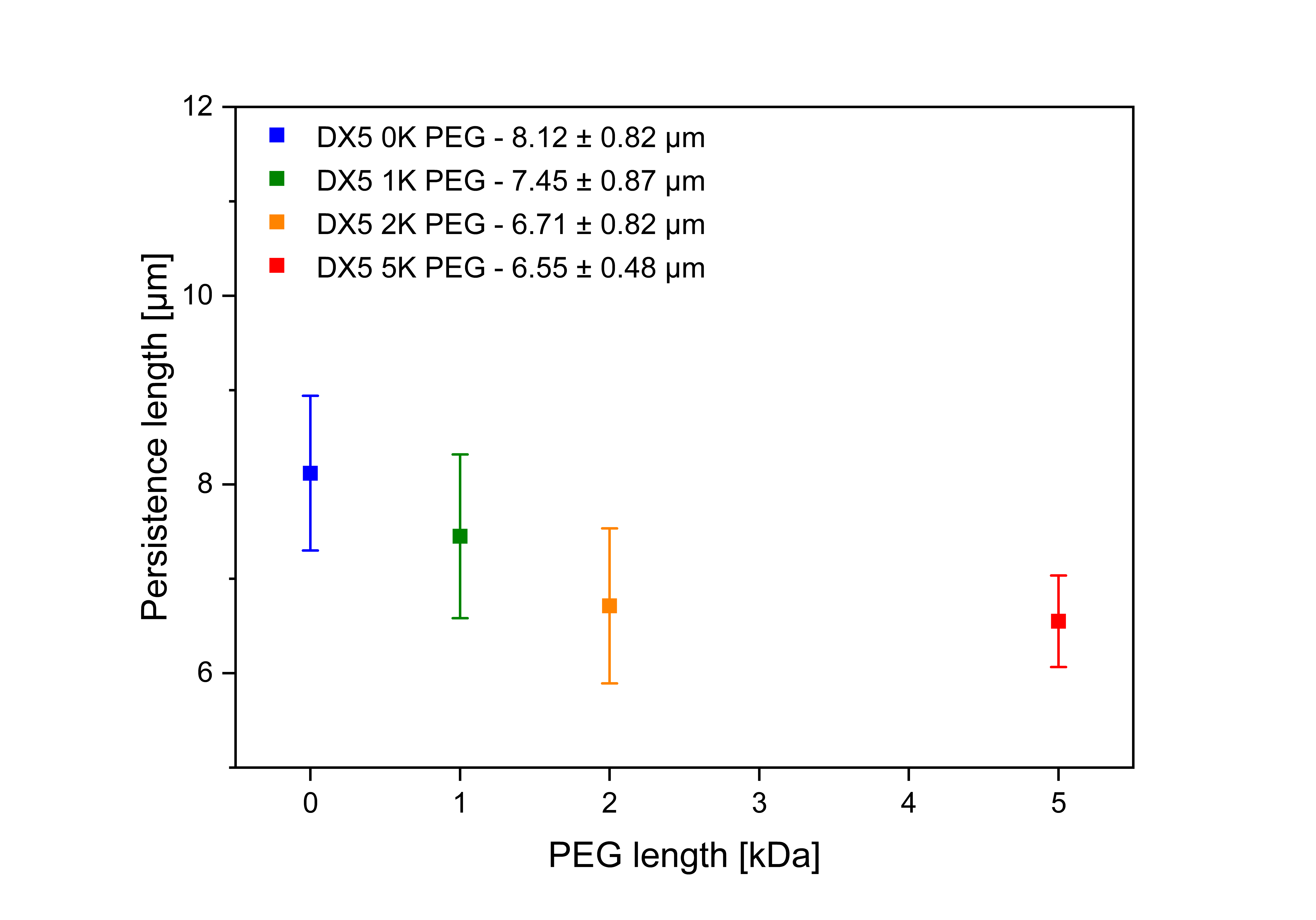

*Figure 4*: Contour lengths and persistence lengths for the hybridized DX5 nanotubes and their PEG modified versions.

**The glassy wormlike chain model.**

The GWLC model used for this study has been deployed and described in earlier studies^1–3^. The GWLC is an extension of the wormlike chain (WLC) for semiflexible polymer networks. Glassy interactions of a test chain with its environment are taken into account by stretching the mode relaxation spectrum of the WLC exponentially. The mode relaxation times of all Eigen modes of (half-) wavelength $\lambda_{n}$ = L/n and mode number n for a WLC with persistence length $l_{p}$ and the transverse drag coefficient $\zeta_{\perp}$ are given by

$$\tau_{n}^{WLC}= \zeta_{\perp} / (\frac{l_{p} k_{B} T \pi^{4}}{\lambda_{n}^{4}}+f \pi^{2}/\lambda_{n}^{2})$$

The relaxation times of the GWLC are modified according to

$$\tau_{n}^{GWLC}= \left\{ \begin{aligned} \tau_{n}^{WLC} if \lambda_{n}\leq\Lambda\\ \tau_{n}^{WLC}e^{{\varepsilon N}_{n}} if \lambda_{n}>\Lambda, \end{aligned} \right.$$

where $N_{n}$ = $\lambda_{n}$/ Λ − 1 is the number of interactions per length $\lambda_{n}$. The average distance between interaction points is expressed in the parameter Λ, L gives the contour length of the test filament and the stretching parameter ε controlls how strong the modes are slowed down by interactions with its environment. $f$ describes a homogeneous backbone tension accounting for existing pre-stress. The complex linear shear modulus in the high frequency regime is given by

$$G^{*}\left( \omega\right)= \Lambda/(5\xi^{2}\chi(\omega))$$

where $\xi$ is the meshsize of the network. The micro-rheological, linear response function $\chi(\omega)$ to a point force at the ends of the GWLC is calculated as

$$\chi\left( \omega\right)=\frac{L^{4}}{\pi^{4} l_{p}^{2} k_{B} T} \sum_{n=1}^{\infty} \frac{1}{(n^{4}+n^{2}f/f_{E})(1+i\omega\tau_{n}^{GWLC}/2)}$$

with the Euler buckling force $f_{E}=l_{p}k_{B}T\pi^{2}/L^{2}$. For the linear regime $f$ is set to zero.

For a more detailed description we would like to refer to Kroy, K. & Glaser, J.^4^

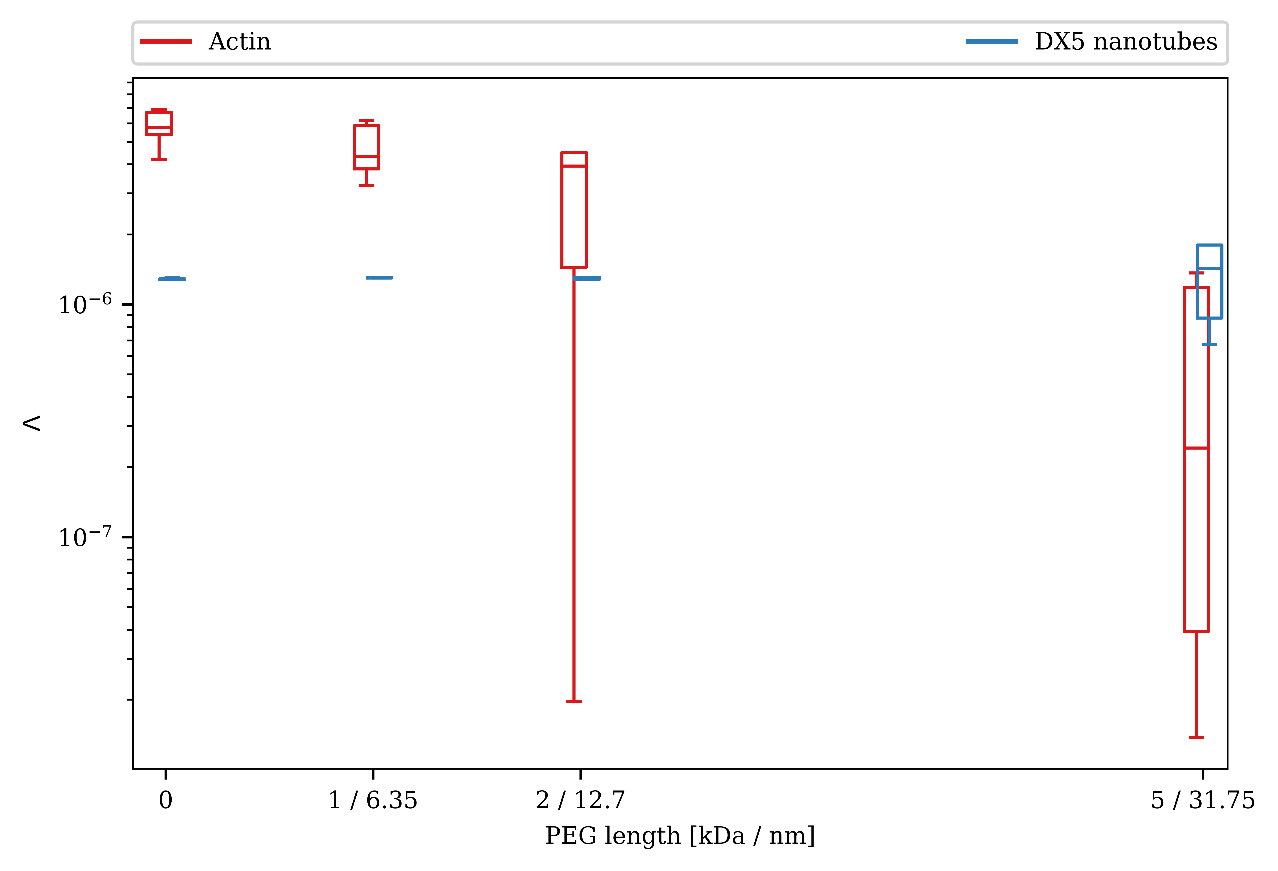

*Figure 5:* Λ values obtained from fitting the experimental data to the GWLC model. Λ is slightly higher as the mesh size for both investigated unmodified systems. While for the DNA based DX5 nanotube system, the value shows only little deviations upon modification with PEG molecules, the actin modifications exhibit a successively decrease of Λ for increasing lengths of PEG spacers attached to filament surfaces.
